# OGP: A Repository of Experimentally Characterized O-Glycoproteins to Facilitate Studies on O-Glycosylation

**DOI:** 10.1101/2020.03.03.975755

**Authors:** Jiang-Ming Huang, Meng-Xi Wu, Yang Zhang, Si-Yuan Kong, Ming-Qi Liu, Bi-Yun Jiang, Peng-Yuan Yang, Wei-Qian Cao

## Abstract

Numerous studies on cancer, biopharmaceuticals, and clinical trials have necessitated comprehensive and precise analysis of protein O-glycosylation. However, the lack of updated and convenient databases deters the storage and utilization of emerging O-glycoprotein data. To resolve this issue, an O-glycoprotein repository named OGP was established in this work. It was constructed with a collection of O-glycoprotein data from different sources. OGP contains 9354 O-glycosylation sites and 11,633 site-specific O-glycans mapping to 2133 O-glycoproteins, and it is the largest O-glycoprotein repository thus far. Based on the recorded O-glycosites, an O-glycosylation site prediction tool was developed. Moreover, an OGP-backed website is already available (http://www.oglyp.org/). The website comprises four specially designed and user-friendly modules: Statistic Analysis, Database Search, Site Prediction, and Data Submit. The first version of OGP repository and the website allow users to obtain vast O-glycoprotein related information, such as protein accession numbers, glycopeptides, site-specific glycan structures, experimental methods, and potential glycosylation sites. O-glycosylation data mining can be performed efficiently on this website, which can greatly facilitates O-glycosylation studies.

## Introduction

Comprehensive and precise analysis of O-glycoprotein would potentially further the current understanding of their roles in many physiological and pathological phenomena, such as intercellular communication [1], hereditary disorders, immune deficiencies, and cancer [2–4]. Great efforts have been made to analyse the complexity of O-glycosylation. And the recent technological advancements in many fields, especially mass spectrometry, generated impressive data on O-glycoproteins [5–11]. However, the lack of up-to-date and curated databases still hindered the archive, query, and utilization of emerging O-glycoprotein data.

Numerous studies have been performed to develop glycosylation-related databases [12–25]. However, most of these databases are focused on N-glycoproteins. Only a few databases contain data on O-glycoproteins. The most extensively used repository, UniCarbKB [13] provides massive N-glycoprotein data mainly and limited O-glycoproteins records. dbPTM [15, 16] is an integrated resource for post-translational modifications (PTMs) containing over 130 types of PTMs; however, it does not provide information regarding site-specific O-glycosylation. O-GLYCBASE [12] provides information regarding both glycans and glycosylation sites and is the most widely used database in O-glycosylation sttudies; however, it has not been updated since 2002. Moreover, it contains merely 189 O-glycoproteins and 2142 O-glycosylation sites, lagging in current O-glycoproteome data. In short, current O-glycoprotein databases are less satisfactory with notable issues including insufficient records, unknown data confidence, outdated or user-unfriendly (Supplementary Table 1).

It can be said that the dearth of O-glycoprotein databases has greatly impeded the development of O-glycosylation study. Recently, large-scale analysis of O-glycosylation sites and intact O-glycopeptides have gradually become possible. For example, Steentofh et al. developed a glycol-engineering method termed “SimpleCell” for large-scale identification of O-glycosylated sites [5]. Yang et al developed a method called “EXoO” for large-scale analysis of intact O-glycopeptides [7]. However, functional studies of O-glycoproteins are yet limited. In addition to the complexity of O-glycosylation, another primary factor limiting studies on O-glycosylation is the difficulty in retrieving information from large data to select candidate O-glycoproteins. Thus, an updated O-glycosylation database providing curated information of protein O-glycosylation status, site-specific O-glycans, analytical methods, and other related information is largely in need and would accelerate studies on O-glycosylation.

In this study, an O-glycoprotein repository named OGP was constructed. OGP contains 9354 O-glycosylation sites and 11,633 site-specific O-glycans mapping to 2133 O-glycoproteins. To our knowledge, OGP is the most comprehensive experimentally characterized O-glycoprotein repository thus far. An O-glycosylation site prediction tool was also developed based on the recorded sites. An OGP-backed website is well established (http://www.oglyp.org/) to facilitate database access. The website contains four modules: Statistic Analysis, Database Search, Site Prediction, and Data Submit. All the aforementioned O-glycoprotein data can be easily obtained on the website. Such a comprehensive, user-friendly, and open-access O-glycoprotein repository would greatly benefit researches on O-glycosylation, development of O-glycoprotein drugs, or clinical studies.

## Construction of the OGP repository

The OGP knowledgebase was constructed by integrating experimentally verified O-glycoproteins between 1998 and 2018 and other existing O-glycoprotein databases [12] (Fig. 1A). All proteins were manually curated, aligned with UniProt entries, and merged. Detailed extraction methods from the literature are described in Supplementary File S1. In total, 9354 glycosylation sites and 11,633 site-specific glycans mapping to 2133 glycoproteins of different species have been recorded in the database (Fig. 1B, C). The scale of the OGP repository is more than 20-fold bigger than the existing O-GlycBase 6.0, which is the largest O-glycoprotein repository thus far (Fig. 1D, E). This database is intended to be further updated to maintain its scale and content by expanding it with more published data.

**Figure 1.**
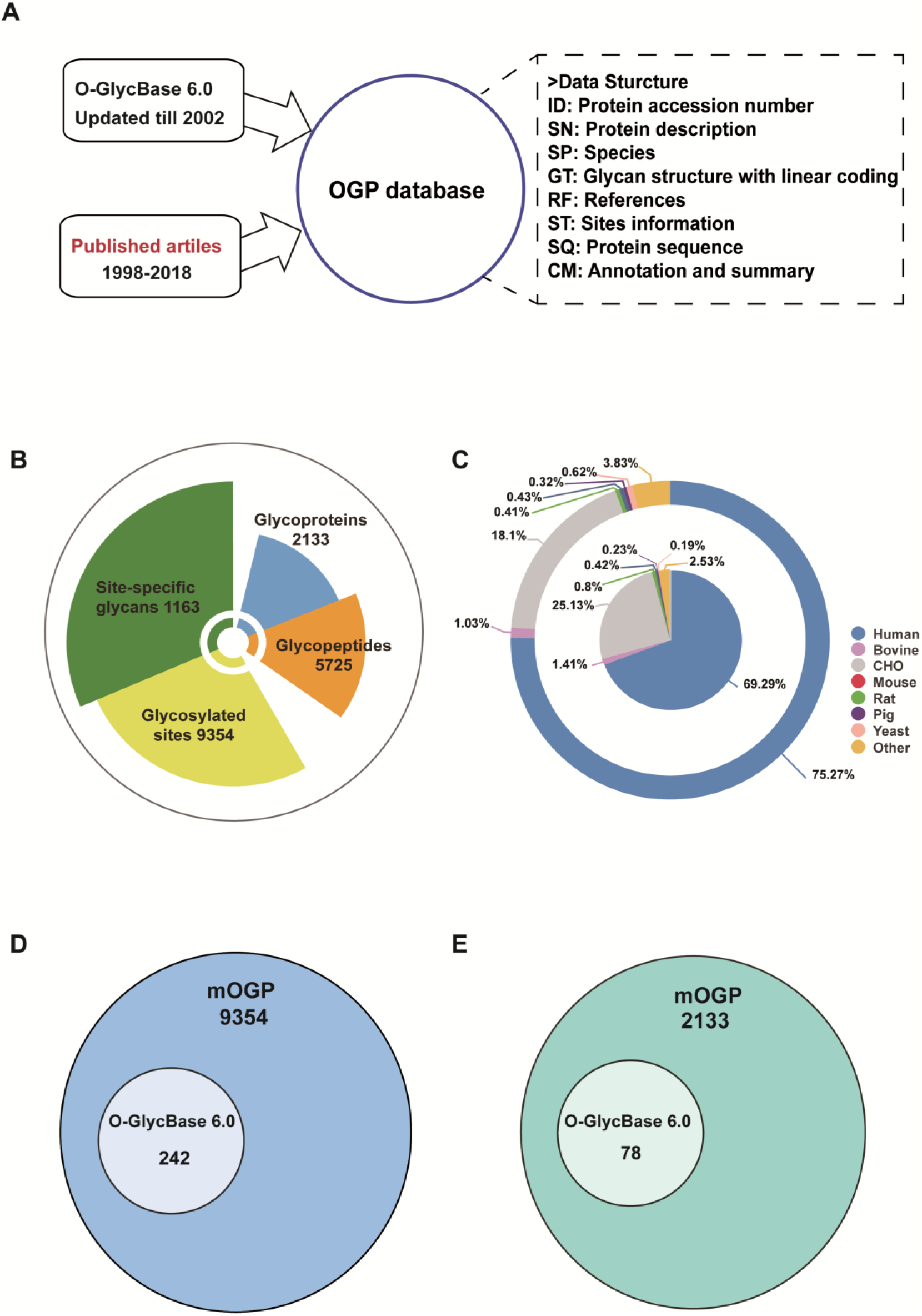
Overview of OGP repository. **A.** OGP Data collection. **B.** The scale of the OGP repository. **C.** Species distribution of O-glycoproteins in OGP. **D.** Comparison of OGP with O-GlycBase 6.0 in glycosylation site level. **E.** Comparison of OGP with O-GlycBase 6.0 in glycoprotein level.

The database records data such as proteins, peptide sequences, glycosylation sites, and site-specific glycans. For each site and site-specific glycan, detailed analytical strategies, such as sample sources, digestion enzymes, enrichment methods, analytical methods, and related studies are integrated. Besides, all glycoproteins recorded in the database have been aligned with their UniProt entries. Thus, additional data including protein sequence annotation, subcellular locations, and other post-translational modifications can be conveniently obtained. To better obtain topological information regarding glycans, a linear coding method (Supplementary File S2) has been used in this database to record site-specific glycan structures. Furthermore, experimental strategies for every glycopeptide, such as immunoprecipitation, gel filtration, and mass spectrometry, were manually extracted, verified, and recorded in the database. These data are easily retrievable from the OGP-backed website.

## Development of an O-glycosylation site prediction model

Since O-glycosylation is highly complex but important, it is significant to better understand its modification pattern [26–29]. As a meaningful trial and for better utilization of OGP, an O-glycosylation site prediction model was developed using meticulously selected O-glycosylation sites with high confidence, recorded in OGP, which were identified at least one solid method to confirm its reliability and unambiguousness; for example, sites validated by Edman Sequencing or HCD and ETD tandem MS. The site prediction model was generated through three primary steps (Fig. 2A, Supplementary File S3): 1) construction of a dedicated training set; 2) optimization of parameters; 3) evaluation of site prediction performance. Through systematic optimization, a dedicated training dataset was established with a 1:1 positive: negative instances ratio for random sampling (1754 positive site-central sequences and 1754 negative site sequences) (Fig. 2B, Supplementary File S3-1.1). Sequences with 11 amino residues were considered preferable (Fig. 2C, Supplementary File S3-1.2). Thereafter, we compared the performance of different algorithms to predict O-glycosylation sites using Weka 3.8 as the data mining tool. The Random Forest algorithm displayed the best performance (Fig. 2D, E, Supplementary File S3-1.3). Finally, and was used to construct the prediction model.

**Figure 2.**
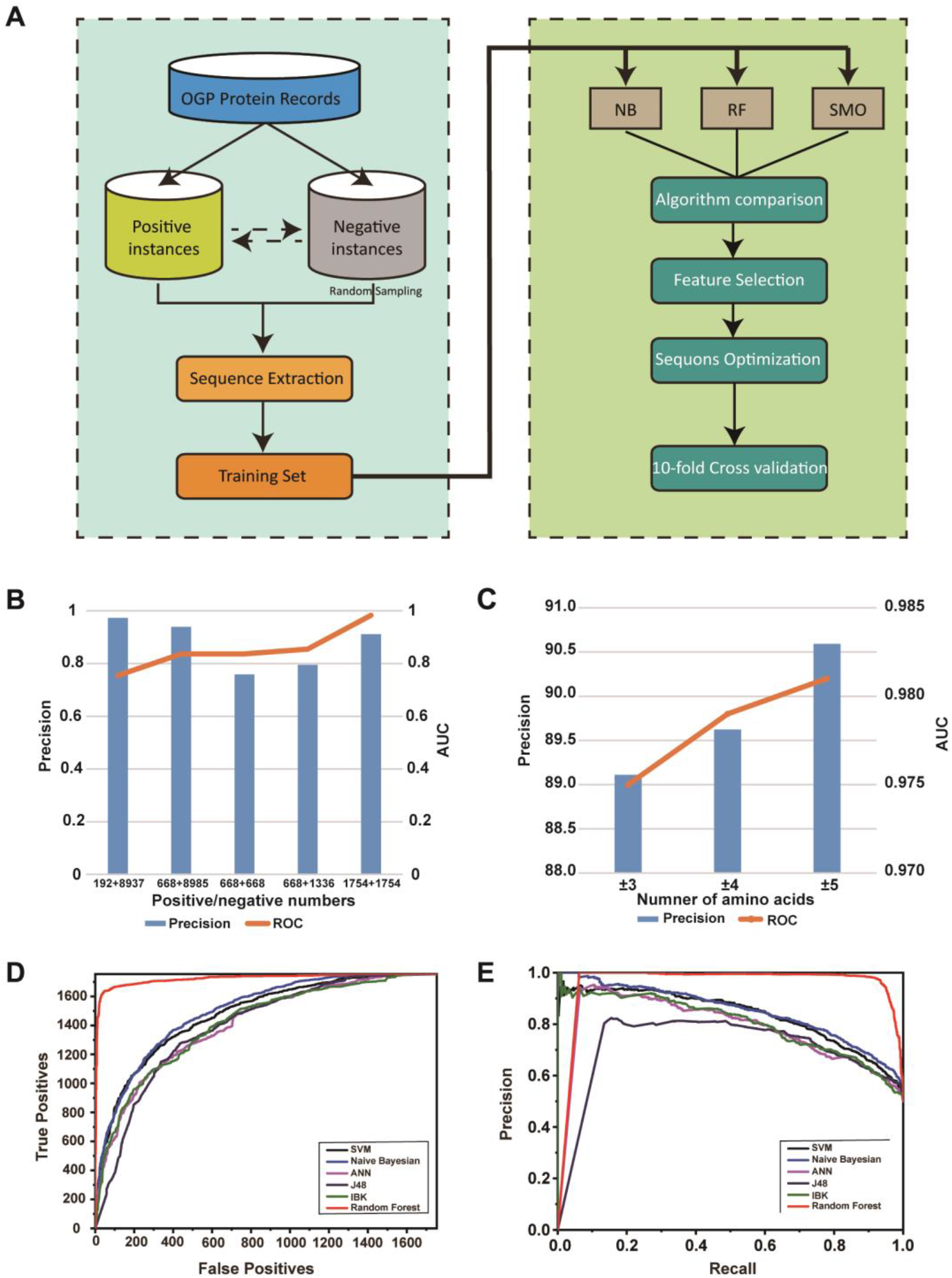
Development of O-glycosylation site prediction model. **A.** Workflow for building OGP based O-glycosite prediction model. **B.** Effect of positives and negatives scales and ratios on model prediction performance (AUC represented for area under curve). **C.** Influence of amino acid residuals on the performance of site prediction model. **D.** ROC curves of each classification algorithm. **E.** PR curves of each classification algorithm.

A ten-fold cross validation was performed and results indicated that the prediction model has high accuracy and sensitivity (ROC value=0.982, Recall value=0.902) (Fig. 2D, E, Supplementary File S3-1.3).

## Construction of the OGP-backed website

Based on the present OGP knowledge base, a dedicated website was constructed using hypertext markup language (HTML), cascading style sheet (CSS), and professional hypertext preprocessor(PHP); The design of the website is shown in Figure 3A. It contains three repositories in the underlying database layer: OGP, Prediction model, and Data submission. OGP is the core database that stores O-glycosylated protein sequences, sites, site-specific glycans, corresponding experimental data, and bibliography; The prediction model contained a model file and an inherent training set; Data submission is design to preserve user-uploaded information. By performing a set of actions including protein query and prediction model training and application in the operation layer, the website outputs four modules: Statistic analysis, Database search, Site prediction, and Data submit. The website is supported by most common web browsers such as Internet Explorer, Mozilla Firefox, Google Chrome, Safari and Opera.

**Figure 3.**
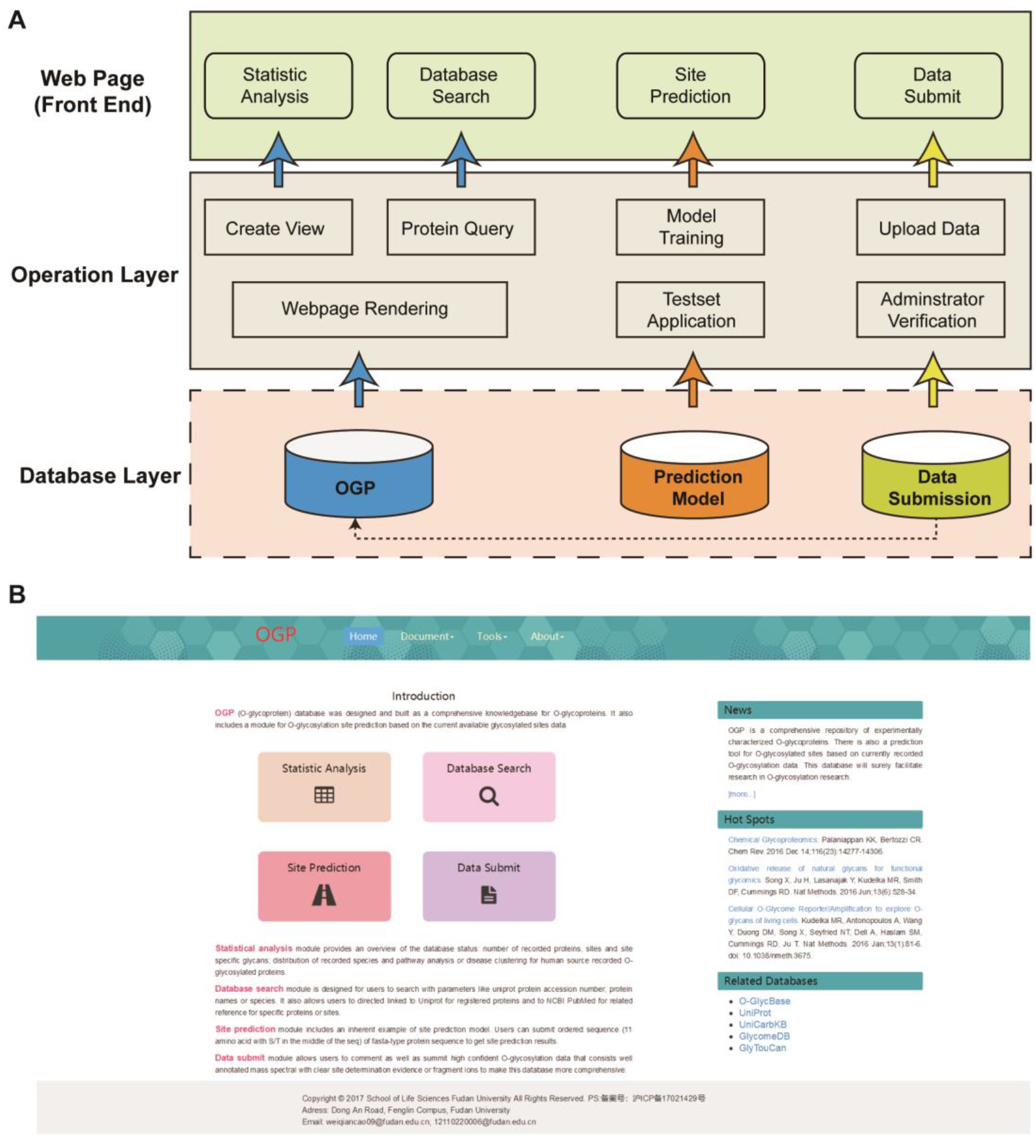
Construction of OGP-based website. **A.** The website structural design. **B.** Homepage of the website.

## Utility and the interface of the OGP website

The OGP-backed website, equipped with a user-friendly graphical interface, is already available (http://www.oglyp.org/) and comprises four main modules: Statistic analysis, Database search, Site Prediction, and Data Submit. Furthermore, other functions including database download, latest studies on O-glycosylation and useful links to databases (UniProt, UniCarbKB, and O-GlycBase) are provided and updated to provide more information for O-glycosylation study. The homepage of this website is shown in Figure 3B. Furthermore, the website provides detailed instructions and frequently asked questions (FAQ) to facilitate users.

The Statistic Analysis module provides an overview of the OGP repository, including the scale of total proteins, glycosylation sites, and site-specific glycans (Supplementary figure 1A), frequency distribution of O-glycoproteins in major taxonomic categories (Supplementary figure 1B), database scale comparison between the OGP and the other primary databases (Supplementary figure 1C), data regarding glycoprotein-involved pathways, molecular and cellular functions of proteins, and related diseases determined through Ingenuity Pathway Analysis (Supplementary figure 1D, E, F). Furthermore, this module provides more essential information. For example, more than 95% of the reported O-glycoproteins are present in mammalians, 55% of which are present in *Homo sapiens*, indicating that O-glycosylation in other species warrants further analysis. All information would be updated in real time with the expansion of the OGP database.

In the Database Search module, users can retrieve O-glycoproteins flexibly by specifying the gene name, protein name, UniProt accession number, or glycan structure (Supplementary figure 2). Figure 4 shows a webpage returning results from a query of fibrinogen gamma chain as an example. These results comprise well-structured data on protein O-glycosylation including basic protein information (*i.e.*, protein name, UniProt accession number, and species, [Figure 4A]), a global map of all recorded O-glycosylation sites of the protein (Figure 4B), experimentally verified O-glycopeptides and site-specific glycans (Figure 4C), and corresponding experimental methods, identifiers and source references (Figure 4D, E).

**Figure 4.**
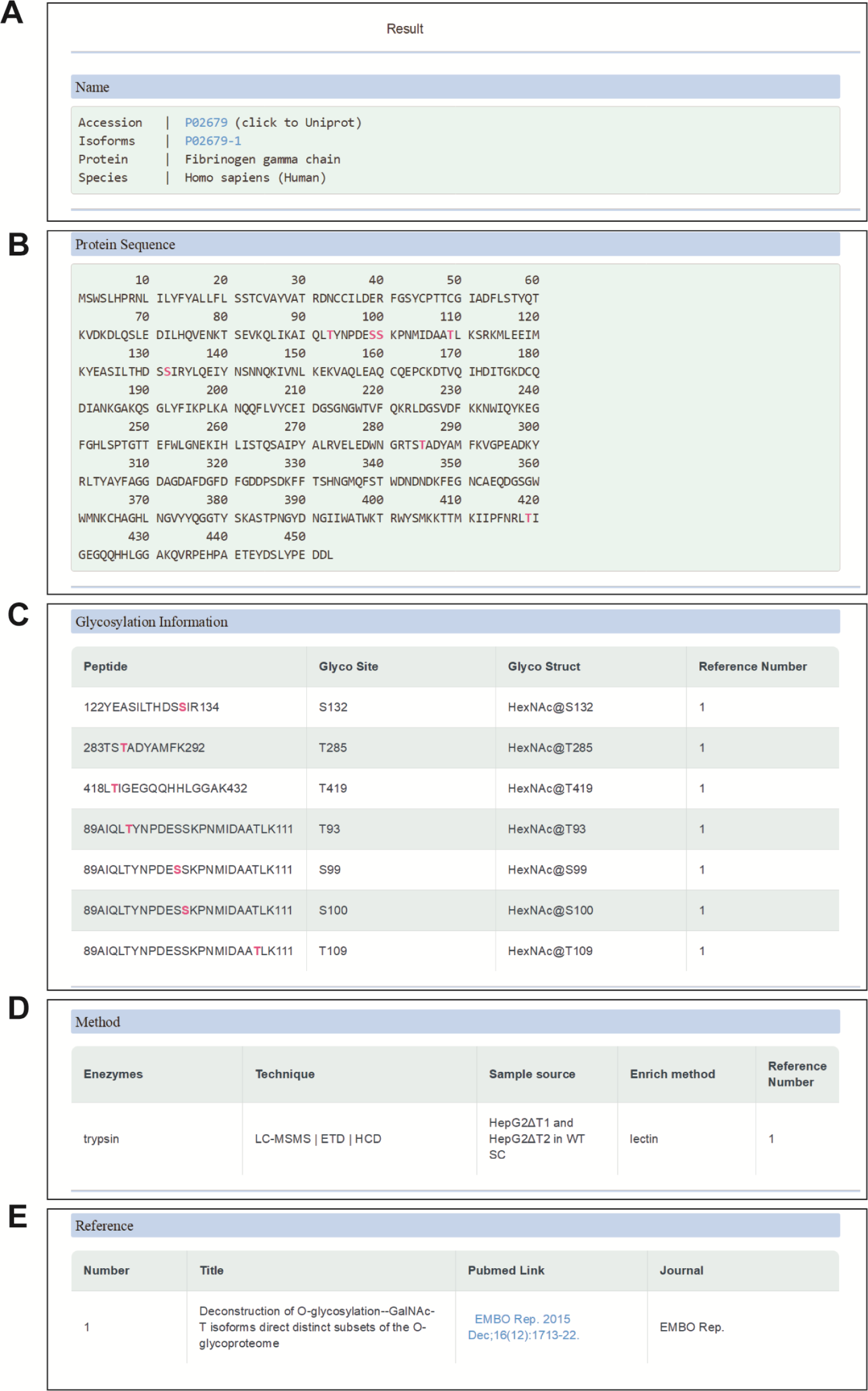
A webpage returning results from a query for Fibrinogen gamma chain. **A.** O-glycoprotein basic information. **B.** Protein Sequences and all recorded O-glycosylation sites with pink highlight. **C**. Experimentally verified O-glycopeptides and site-specific glycans. **D.** Corresponding experimental methods. D. Related source references.

The site prediction model developed herein has also been incorporated into the website to enable O-glycosylation site prediction. As is shown in Supplementary figure 3A, users can either fill out the template file with aligned site-centered sequences as instructed or simply upload a typical protein FASTA-type file and click on “predict” (Supplementary figure 3A). The prediction results for each site can be then displayed directly on the right side of the web page. Prediction scores range between 0 and 1; scores higher than 0.5 indicate positive sites, while those less than or equal to 0.5 indicate a highly probably non-O-glycosylated site. The higher the score, the greater the probability of a site being O-glycosylated and *vice versa*. The results can also be downloaded as shown in Supplementary figure 3B.

The Data Submit module enables users to upload new data into the OGP database and insert comments. All the new submitted data and comments are carefully recorded in a backend database and revised manually by experts at regular intervals. Both a template form and an online form are accepted during a submission. What’s more, when users upload the data by file, there will be a real time feedback shown below to inform users of those O-glycoproteins already in OGP database.

In addition, the database is accesible from OGP website. Download page can be found in the drop-down menu of tools on OGP homepage. The detailed top 500 entries could be directly downloaded (http://www.oglyp.org/download.php). Besides, there is a basic version of database, which provides all the O-glycoprotein Accessions and their corresponding O-glycosylation sites, for users to download freely (http://www.oglyp.org/download.php). The whole database could also be provided if users apply for it through E-mail request. The application guide is illustrated on the website (http://www.oglyp.org/download.php).

## Conclusion and Discussion

The OGP repository, containing 9354 O-glycosylation sites and 11,633 site-specific O-glycans mapping to 2133 O-glycoproteins, is the most comprehensive, experimentally verified O-glycoprotein repository thus far. All data contained in the OGP repository have been manually curated, and protein sequences have been aligned with UniProt entries and merged. Based on recorded site data, an O-glycosylation site prediction tool has been developed to facilitate the prediction of O-glycosylation sites. The OGP-based website is already available (http://www.oglyp.org/) and contains four specially designed, user-friendly, functional modules: Statistic Analysis, Database Search, Site Prediction, and Data Submit. The initial version of the OGP repository and OGP-based website provides various data on O-glycoproteins, such as protein access number, sequence, site-specific glycan structures, experimental methods, and potential glycosylation sites. Mining of data on O-glycosylation can be carried out efficiently using this website. The OGP repository would greatly facilitate studies on O-glycosylation. The scale and content of this database is intended to be continuously expanded in subsequent versions of the OGP repository.

## Availability

OGP prediction tool is freely available at http://www.oglyp.org/predict.php. OGP database is freely available at http://www.oglyp.org/download.php.

## Authors’ contributions

HJ developed the site prediction tool, participated in the database construction and helped to draft the manuscript. WM performed the database uniform, participated in the database and website construction and helped to draft the manuscript. ZY participated in website construction. KS participated in data collection. LM helped to revise the manuscript. JB participated in data collection. YP conceived the study, helped to draft the manuscript. CW designed the database, conceived the study, and draft the manuscript. All authors read and approved the final manuscript.

## Competing interests

The authors have declared no competing interests.

## Acknowledgements

This work was supported by grants from the National Key Research and Development Program (2018YFC0910300, 2016YFA0501303 and 2016YFB0201702), the National Natural Science Foundation of China Project (91853102, 31600665 and 31570825). We would like to thank Mr. Jian-Qiang Wu to help with website construction, and Elsevier Premium Language Editing Services to help with the language editing.

## Notes

http://www.oglyp.org/

