## Supplementary material for "OGP: A Repository of Experimentally Characterized O-Glycoproteins to Facilitate Studies on O-Glycosylation": SI

### Supplementary File S1 Literature extraction

Literatures published between 1998 and 2018 that matched searching key words of "O-glyc*" and "mass spect*" was retrieved from Web of Science core collection. A total of 1775 records were carefully reviewed manually. Relative strict criteria for O-glycopeptide selection were applied when screening: identification must include at least one solid validation method to confirm its reliability and unambiguousness; for example, sites validated by Edman Sequencing [1] and HCD, ETD tandem MS [2]; glycan structures verified by lectin affinity chromatography [3] or sequential enzymatic released techniques [4] to fully elucidate the glycan structures. Identification using merely MS1 m/z putative deduction or single broad-specificity lectin affinity was discarded. Identified glycoprotein, glycosylation sites and corresponding glycan structures were then filtered and recorded. In addition, sample information like source of species (*i.e.* human, mouse, and bovine), organs (*i.e.* liver, kidney, and plasma), identification and verification techniques (*i.e.* tandem MS, NMR, and lectin affinity chromatography) were recorded at length. Every entry recorded was manually checked and aligned with its UniProt information on both protein accession number, protein name, sequences and positions of glycosylation site to confirm explicit O-glycoprotein information. Links to UniProt were also provided on queried pages to assist cross-reference. What's more, references for every single glycopeptide identification were listed with article titles, pubmed-formatted citations, publication date, and its links to Pubmed.

### Supplementary File S2 Linear coding for glycan

To facilitate record, storage and spread, all branched glycans were transformed to linear-coded structure forms [5] in OGP to preserve the structure information at greatest extent, as depicted in Figure S1. To be concrete, mass spectrometry identified glycan units GalNAc, Gal, NeuAc and NeuGc were unified as HexNAc, Hex, NeuAC and NeuGc respectively due to the inability to differentiate real saccharide on merely m/z. Branched structures were transformed to nested forms with baskets adjacent to branched nodes (left). Linkages or bond information like alpha, beta conformation and/or 1,3-, 2,4- bond linkages, were also manually checked and recorded, although only a few of glycans have that information. Furthermore, glycans were assigned to corresponding sites with a connector @, followed by abbreviated threonine, serine or tyrosine and its absolute position within the glycoprotein. Different glycans for each glycosylation site or from the same glycopeptide groups (glycopeptides bearing the same glycosylation sites as generated from different proteases) were also collected, recorded and separated using semicolons. As a result, variant glycan compositions on a single site can well demonstrate the overall understanding and the heterogeneity of O-glycosylation at present and thus can serve as useful information for researchers.

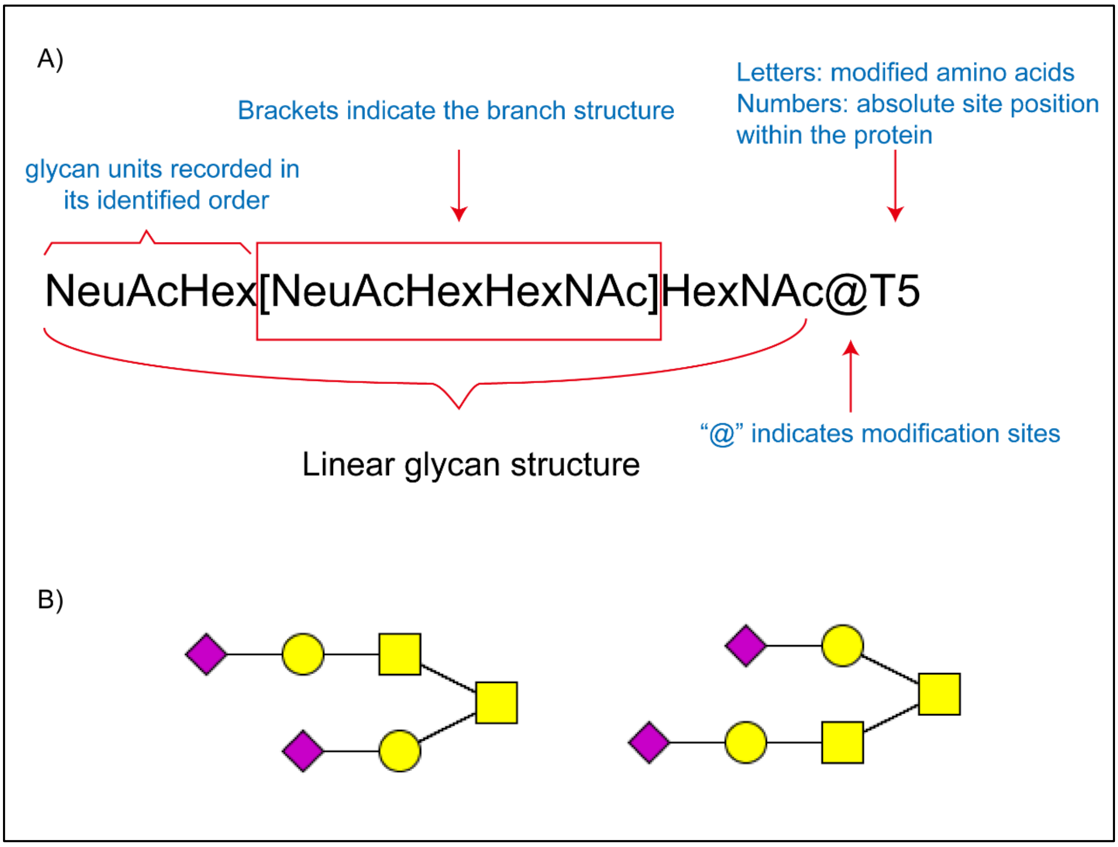


Figure S1. A) linear glycan structure recorded in OGP and instructions; B) putative glycan structure of corresponding linear structure in A).

### Supplementary File S3 Construction of OGP-based site prediction model

The process of building OGP based O-glycosylation site prediction model mainly contained three steps, as is shown in Figure 2A. Firstly, a dedicated training set was constructed with positive O-glycosylation sties and negative O-glycosylation sites. The positive O-glycosylation sites were manually extracted from experimentally identified and verified human O-glycosylated sites from OGP. Negative sites were randomly generated from the residual sequences of the corresponding protein in UniProt (downloaded by the date of 2016.06.27) (Step 1). Secondly, a series of parameter optimization process were conducted to improve the performance of the prediction model and maximally avoid oversampling effects. The amino acid length as well as the optimized classification methods were tested to figure out the best algorithms for prediction (Step 2). Finally evaluation of site prediction performance was conducted with 10-fold cross validation (Step 3) to further prove the accuracy and stability of this model.

#### 1.1 Optimization of positive/negative instances ratio

We firstly inspected impact of positives and negatives proportion on model prediction performance. WEKA 3.8 and random forest algorithms were chosen as preset condition for O-glycosylated site prediction. Briefly, human O-glycosylated sites recorded in OGP were manually extracted as positive instances, while non-reported and non-experimental verified serine/threonine sites on O-glycoproteins recorded in OGP database were pretreated as negative sites (downloaded from UniProt Database at 2016.06.27). Since the crude training set was highly unbalanced (positive/negative (192/8937) ≈ 1/16), high average classification accuracy but poor positive site classification accuracy was observed on the crude training set. As is shown in the confusion matrix in Table S1, the prediction precision for positive instances was only 2.127% (4/188), though the average precision is as high as 97.8928%. To solve this problem, we tried both increasing positive instances from new records, such as multiply positive samples (or so-called oversampling), and decreasing the proportion of negative instances (down-sampling) respectively. As is shown in Figure 2B and Table S1, incorporating with more O-glycosylated sites from newly published data promotes the positive prediction precision (4/188, 507/601, 1692/69) while lowering the ratio of negative to positive instances promote the ROC curve area of the classification model, which increased the robustness of prediction models. However, considering that simply multiplied positives greatly increased the risk of over fitting (namely assigning same modification site patterns) and was prone to larger generalization error, we finally adopted down sampling methods that can also improve glycosylated site prediction precision as well as kept the stability of prediction model. As a result, a total of 1754 positive site-central sequences and 1754 negative site sequences were included as final training set

It's worth mentioned that the prediction performance can still be improved with the extension of O-glycosylation records in the future to serve as a more accurate prediction tool.

**Table S1 Effect of scale and ratio of positives and negatives on model prediction**

| TP ratio | Sampling | Precision | Confusion matrix# | |
|---|---|---|---|---|
| | | | TP/FP | TN/FN |
| 192+8937 | Early version | 97.8928% | 4/188 | 8869/68 |
| 668+8985 | Median version | 94.1670% | 121/547 | 8968/16 |
| 668+668 | 1:1 down sampling | 75.4491% | 507/161 | 501/167 |
| 668+1336 | 1:2 down sampling | 99.3995% | 3340/0 | 8910/74 |
| 1754+1754 | Final version | 90.5929% | 1692/62 | 1473/281 |

#: TP, true positive; FP, False Positive; TN, True Negative; FN, False Negative.

#### 1.2 Optimization of sequon length

It was reported that protein O-glycosylation is catalyzed by 20 GalNAc transferase isoenzymes [6], and the enzymatic catalysis mechanism. Therefore, amino acid (AA)

sequon or modified-site-centered AA length was expected to have great impact on the successfulness of a site being O-glycosylated. Sequons of 3, 4 and 5 AAs forward and backward off potential modification sites were extracted to test their effects on prediction model performance. Deficient of AAs was complemented with letter “X”. As is shown in Figure 2C, with the number of AAs increasing, prediction precision and area under curve (AUC) slightly increased, indicating longer sequences had better performance. Taken into consideration both the fact that the catalytic domain of GalNAc transferases was about 12 AAs and the risk of increasing error rate with complement for terminal serines or threonines for longer sequences, we finally chosen ±5 amino acids as a prediction sequence for this tool.

**1.3 Comparison of different algorithm’s performance on site prediction**

Due to the complexity of O-GalNAcylated site catalytic mechanism, a variety of classification algorithms have been tested for O-glycosylated site prediction, including supporting vector machine (SVM), artificial neural networks, random forest and et.al [7]. In this study, we carefully surveyed the performance of different classification algorithms on predicting probability of O-glycosylation to figure out the most suitable one. The modeling tasks were performed mainly on an open-source machine-learning tool Weka 3.8 [8] (Waikato Environment for Knowledge Analysis). Parameters for each classification algorithms was well-optimized and listed as follows:

**Support vector machine (WLSVM):**

weka.classifiers.functions.LibSVM -S 0 -K 2 -D 3 -G 0.0 -R 0.0 -N 0.5 -M 40.0 -C 1.0 -E 0.001 -P 0.1 -B -seed 1

with classification type set as C-SVC and kernel type set as radial basis function: exp(-gama*|u-v|^2)

**Multi-layer neural network (MultilayerPerceptron):**

weka.classifiers.functions.MultilayerPerceptron -L 0.3 -M 0.2 -N 500 -V 0 -S 0 -E 20 -H a -R

with layer number set as ‘a’ = (attributes + classes) / 2.

**C4.5 decision tree(J48):**

weka.classifiers.trees.J48 -C 0.25 -M 2

**KNN(J48)**

weka.classifiers.lazy.IBk -K 5 -W 0 –A

with the number of neighbors(KNN) optimized as 5

**Random Forest**

weka.classifiers.trees.RandomForest -I 500 -K 0 -S 1

with number of tree was set as 500 which was tested to be stable for the classification model in this case

All classification modeling were performed on the same training set mentioned in Supplementary File III-1.1 under a 10 fold cross-validation. The performances of each algorithm were shown in Table S2, Figure 2D and Figure 2E. In summary, random forest outperformed other classification methods with a prediction precision of 90.9% and ROC area of 0.983 and was finally adopted as the final algorithms for O-glycosylation site prediction. It was currently the highest prediction precision model for O-glycosylation reported.

To our understanding, there might be two reasons for the superior performance of random forest over other algorithms: 1) the multi-decision characteristics of random forest better mimic the multiple transferases-regulated O-glycosylation; 2) the site training set adopted in our modeling process is the largest by far so that the modeling can cover more comprehensive site pattern compared with existing methods.

**Table S2 Prediction performance of several algorithms.**

| **Algorithms** | **TP Rate** | **FP Rate** | **Precision** | **Recall** | **F-Measure** | **ROC Area** | **Time Cost(s)** |
|---|---|---|---|---|---|---|---|
| SVM | 0.764 | 0.236 | 0.765 | 0.764 | 0.764 | 0.842 | 7.33 |
| ANN | 0.725 | 0.275 | 0.726 | 0.725 | 0.725 | 0.804 | 5355.3 |
| Naïve Bayesians | 0.771 | 0.229 | 0.772 | 0.771 | 0.77 | 0.856 | 0.01 |
| J48 | 0.734 | 0.266 | 0.734 | 0.734 | 0.734 | 0.795 | 0.05 |

| | | | | | | | |
|---|---|---|---|---|---|---|---|
| Ibk | 0.724 | 0.276 | 0.732 | 0.724 | 0.722 | 0.804 | 0.01 |
| Random Forest | 0.909 | 0.091 | 0.915 | 0.909 | 0.909 | 0.983 | 1.79 |

### Supplementary Figure

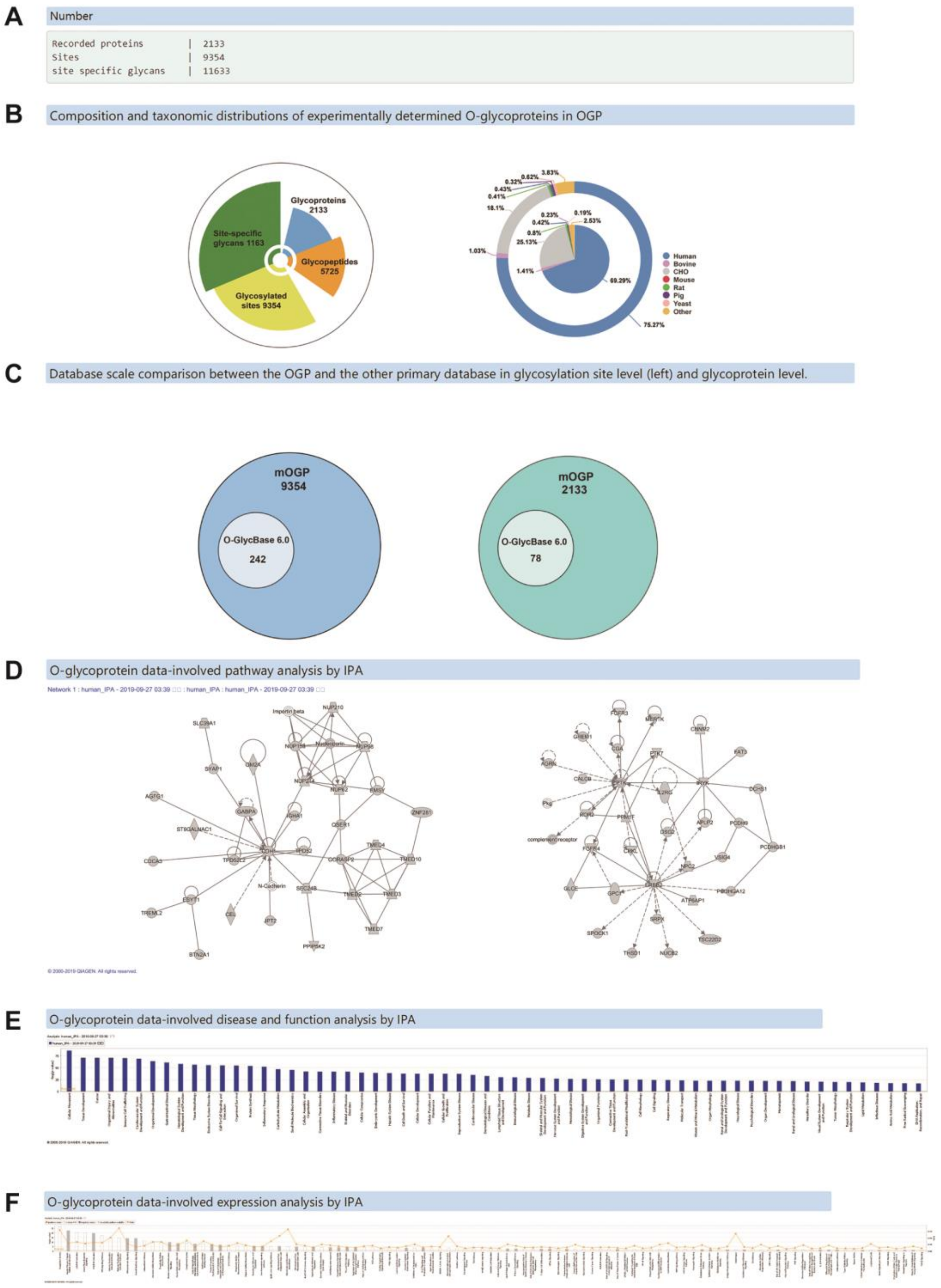


**Supplementary figure 1  Demonstration of statistical analysis module in OGP website.**

**A.** The scale of OGP database. **B.** Composition and taxonomic distribution of O-glycoproteins in OGP database. **C.** Database scale comparison between the OGP and the other primary databases in glycosylation site level (left) and glycoprotein level. **D.**

Data regarding glycoprotein-involved pathways. **E.** Glycoprotein-involved diseases determined through Ingenuity Pathway Analysis. **F.** Molecular and cellular functions of O-glycoproteins in OGP database.

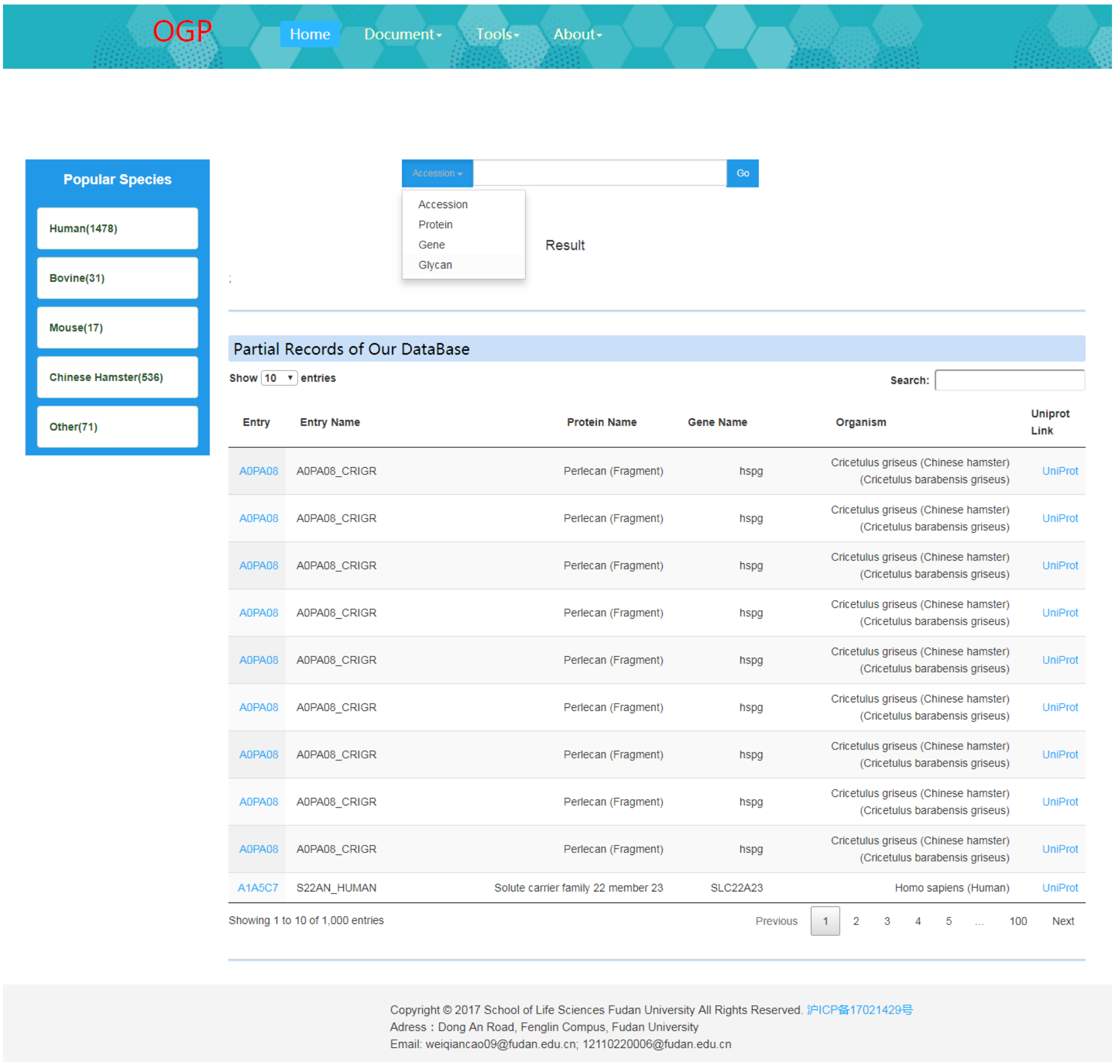


**Supplementary figure 2 The webpage of Database Search Module.**

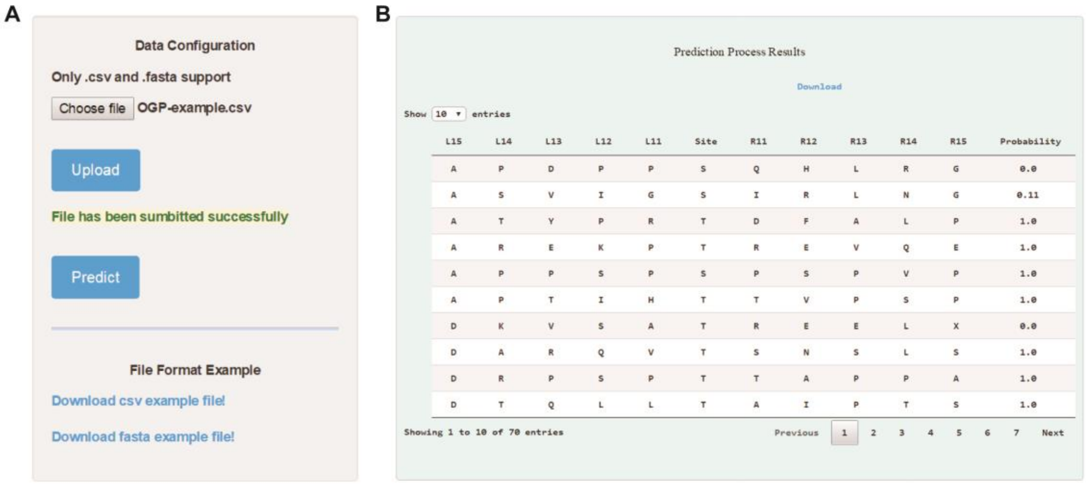


**Supplementary figure 3 The webpage of Site Prediction Model.**

**A.** Submission of required prediction data. **B.** The prediction results for each site

**Supplementary Table 1 Comparison of OGP with other available O-glycoproteome databases.**
